# Time-resolved cryo-EM reveals early ribosome assembly in action

**DOI:** 10.1101/2022.11.12.516267

**Authors:** Bo Qin, Simon M. Lauer, Annika Balke, Carlos H. Vieira-Vieira, Jörg Bürger, Thorsten Mielke, Matthias Selbach, Patrick Scheerer, Christian M. T. Spahn, Rainer Nikolay

## Abstract

Ribosome biogenesis is a fundamental multi-step cellular process in all domains of life that involves the production, processing, folding, and modification of ribosomal RNAs (rRNAs) and ribosomal proteins. To obtain insights into the still unexplored early assembly phase of the bacterial 50S subunit, we exploited a minimal *in vitro* reconstitution system using purified ribosomal components and scalable reaction conditions. Time-limited assembly assays combined with cryo-EM analysis visualizes the structurally complex assembly pathway starting with a particle consisting of ordered density for only ∼500 nucleotides of 23S rRNA domain I and three ribosomal proteins. In addition, our structural analysis reveals that early 50S assembly occurs in a domain-wise fashion, while late 50S assembly proceeds incrementally. Furthermore, we find that both ribosomal proteins and folded rRNA helices, occupying surface exposed regions on pre-50S particles, induce, or stabilize rRNA folds within adjacent regions, thereby creating cooperativity.

## Introduction

The assembly of ribosomal subunits, the largest ribonucleoprotein particles in the cell, relies on the intrinsic affinities of their rRNA and protein components that interact in a hierarchical manner ^1,2^, involving protein-induced conformational changes in rRNA ^3^. The resulting cooperativity of assembly ensures the completion of ribosomal subunits ^4^. The bacterial 50S large ribosomal subunit is composed of 33 proteins (L-proteins), one 23S and one 5S rRNA. The 23S rRNA consists of six architectural domains that, stabilized by the L-proteins, define the shape of the 50S subunit with its three protuberances ^5–7^. Cryo-EM studies of bacterial 50S precursors both purified from cells ^8–13^ and obtained by *in vitro* assembly ^14^ have elucidated the later phase of the maturation pathway.

The most immature 50S precursor states reported contained an already fully formed core, consisting of 23S rRNA domains I, II, III, VI and early binding L-proteins ^9,11,14^. In this core particle about 50% of all protein and RNA residues are already maturely positioned ^14^. Further assembly involves the formation of the L1 stalk, the central protuberance (CP), the GTPase associated center (GAC), and the stalk base. 50S assembly terminates with folding of the peptidyl transferase center (PTC) and the surrounding rRNA elements referred to as functional core (FC) ^9–11,14–17^. The events after structural formation of the large subunit’s core appear to be evolutionary conserved. Hence, similar assembly intermediates have been observed for the eukaryotic 60S subunit ^18–20^ and large subunit precursors derived from mitochondria, which shed light on the latest period of assembly, detailing the finely tuned steps leading to completion of the subunit’s active site ^21–26^, although the involved biogenesis factors differ substantially ^12^.

In comparison, early assembly of the large ribosomal subunit remains elusive from a structural standpoint. Pioneering studies visualizing nucleolar derived samples from yeast provided first structural insight into intermediate states before core formation has been completed ^27,28^. However, large subunit precursors exploring the earliest stages of their assembly escaped structural analyses so far.

## Results

### Time course of 50S *in vitro* reconstitution

To focus on early events of 50S assembly, we adopted our previous approach ^14^ and performed step 1 of the *in vitro* reconstitution assay ^29^ as a time course reaction, analyzed sucrose gradient profiles, translation activity of the assembled particles (after a full step 2 incubation), and determined the protein occupancy, using quantitative mass spectrometry (q-MS) **(Methods, Fig. 1A, and Extended Data Fig. 1A-K)**. Before heat incubation, the sample migrates in a sucrose gradient as a 33S precursor **(Fig. 1B)** with reduced amounts of the L-proteins bL32, uL30, uL14, uL29, bL19, bL28 and uL16 **(Extended Data Fig. 1K)**, and exhibits no translation activity **(Fig. 1C)**, in agreement with previous studies ^30^. As expected, both the portion of faster migrating particles and the translation activity increased with the time of incubation **(Fig. 1C, and Extended Data Fig. 1B-H)**.

**Fig. 1:**
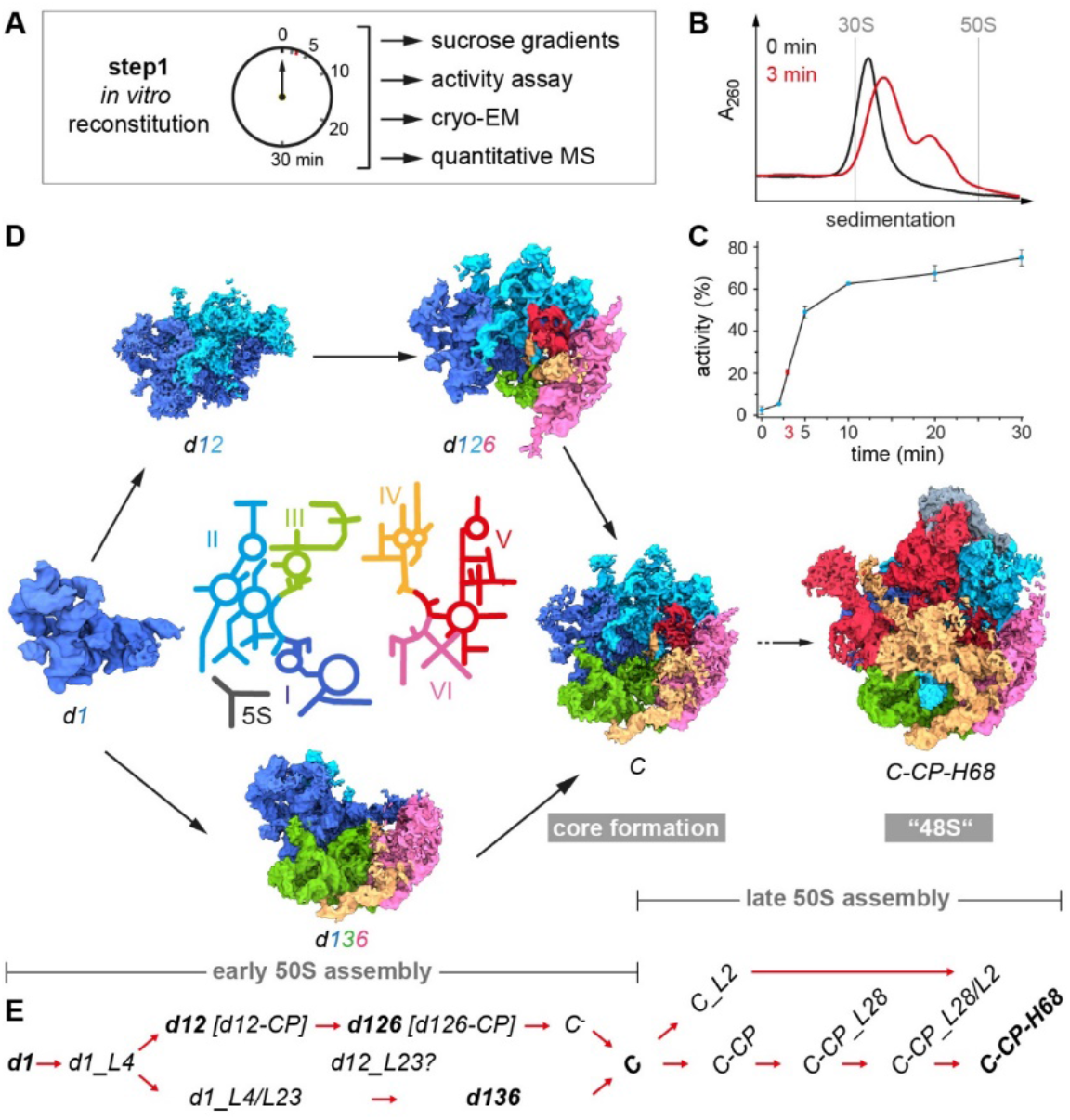
Time course of 50S step 1 *in vitro* reconstitution with structural and biochemical analyses. **A)** Experimental setup. **B)** Sucrose gradient profiles of *in vitro* reconstitution samples incubated for 0 min (black) or 3 min (red), respectively. **C)** *In vitro* translation assay indicating the relative activity of subunits after the indicated time of incubation under step 1 conditions plus subsequent incubation under step 2 conditions for 90 min. **D)** In the center, a color-coded 2D map of the 23S rRNA with the individual domains labeled from I-VI (5S, 5S rRNA). Cryo-EM maps of selected states derived from the step 1 reaction after 3 min of incubation appear in the same color-code. **E)** Nomenclature for states occurring during early and late 50S assembly. States in bold are shown in (D). *d*, 23S rRNA domain; *C*, 50S core; CP, central protuberance; H68, helix 68.

The sucrose gradient profiles revealed that after 3 min of reaction, the sample exhibits its largest complexity with the majority of precursors in the 33S region and a significant amount of faster migrating particles (**Fig. 1B**). Nevertheless, particles in the 3 min sample contain all L-proteins (except bL31 and bL36) with occupancies of >70% **(Extended Data Fig. 1I)**. Hence, we selected this sample for a detailed cryo-EM analysis. After extensive sorting and multi-particle refinement, we were able to disentangle 16 different states, with nominal resolutions between 3.0 and 6.6 Å **(Extended Data Fig. 2 and Extended Data Fig. 3)**. While some of the obtained states are related to the 41S-like and 48S-like particles of the late phase of assembly that we have previously obtained with a full 30 min incubation ^14^, the time-course approach resulted in nine novel states that were clearly less complete than the core particle. Whenever in the following we describe the presence, or appearance of an L-protein, an rRNA element or a domain, it refers to detectable density in the corresponding cryo-EM map that requires rigid incorporation of the element into the main particle.

### Structure-based pathway for early assembly

The earliest state we identified contained strong density for ∼500 bp of domain I and the L-proteins uL22, uL24 and uL29 (referred to as state *d1*). We find that at lower resolution, our particles often exhibit additional density that is difficult to interpret molecularly, but may be caused by elements that are being stably incorporated but are still in a dynamic state. Nevertheless, state *d1* represents the earliest pre-50S intermediate reported so far **(Fig. 1D, and Extended Data Fig. 4A)**. Hence, 50S assembly *in vitro* starts with the stable formation of the 5’end containing domain I, despite the presence of the entire 23S rRNA molecule. Interestingly, uL24, located in the center of *d1*, is found among the first binders **(Extended Data Fig. 1I)** rationalizing its role as assembly initiator ^30^. To better understand the molecular anatomy of the subunit, we arranged selected cryo-EM maps in a logical order and noticed that early assembly can progress at least along two putative routes **(Fig. 1D)**.

Instead of numbering the individual states successively, we designed a more specific nomenclature **(Fig. 1E)**. The early states were designated according to the 23S rRNA domains whose densities they exhibit (e.g., *d1, d12, d126*, etc.). In order to distinguish states very similar in composition, additional structural elements were added to their names (e.g. *d1L4/L23*, densities for uL4 and uL23 etc.). Once densities for domains I, II, III, VI, and their interacting proteins are present, the large subunit’s core has completely formed (state *C*). To differentiate states occurring during late assembly, emerging structural features such as the central protuberance, or helix 68 were added to their names (*C-CP*, or *C-CP-H68*).

While state *d1* can mature to *d12* and *d126*, an alternative route involves state *d136*. Both states *d126* and *d136* can convert into state *C* (*d1236*), a process we refer to as core formation, that enhances base pairing between the 5’ and 3’ ends of 23S rRNA. Consequently, core formation *in vivo* can only be accomplished after transcription of the entire 23S rRNA molecule. Subsequently, state *C* transitions via intermediate steps into state *C-CP-H68* (48S) **(Fig. 1D and E, and Extended Data Fig. 4 and 5)**. Interestingly, some of the 23S rRNA domains indeed appear to be folding units that are intrinsically rigid, but flexible relative to each other **(Movie 1)**. Domains I and VI exhibit full cryo-EM density within their domain boundaries as soon as they become detectable as parts of discrete states (**Fig. 1D;** *d1, d126* and *d136*). Defined regions of domains II and III show density in states *d1_L4* and *d1_L4/L23*, respectively, before subsequent concerted appearance of their remaining densities (**Extended Data Fig. 4)**. In the case of domain II only density for the central region appears and extended parts such as A-site finger and GTPase associated center (GAC) form in late assembly. Taken together, early 50S assembly occurs in a domain-wise fashion, while late 50S assembly proceeds incrementally.

### Detailed molecular morphogenesis - Contribution of L-proteins

L-proteins concentrating on the back site of the subunit are organized in clusters, containing at least one early assembly protein, providing contact points for later binding proteins and connecting adjacent rRNA domains **(Fig. 2A)**. To exemplify in more detail how certain L-proteins contribute to 50S assembly, we compared states *d1, d1_L4* and *d1_L4/L23* **(Fig. 3A-E, and Movie 2)**. State *d1* is lacking density for uL4 **(Fig. 3A)**, while *d1_L4* exhibits densities for uL4 along with domain II helices H27-31 **(Fig. 3B)** Similarly, state *d1* is lacking density for uL23 **(Fig. 3D)**, while *d1_L4/L23* exhibits densities for uL23 along with domain III helices H49-53, including the interjacent bulges **(Fig. 3E)**. Thus, uL4 and uL23 appear to contribute to the formation of local seeds to promote further assembly of domains II and III, respectively **(Fig. 2B-K)**. However, while uL4 is present in all downstream assembly intermediates, presence, or absence of uL23 determines, whether assembly proceeds along route 1-3-6 or 1-2-6 **(Fig. 2E-G)**.

**Fig. 2:**
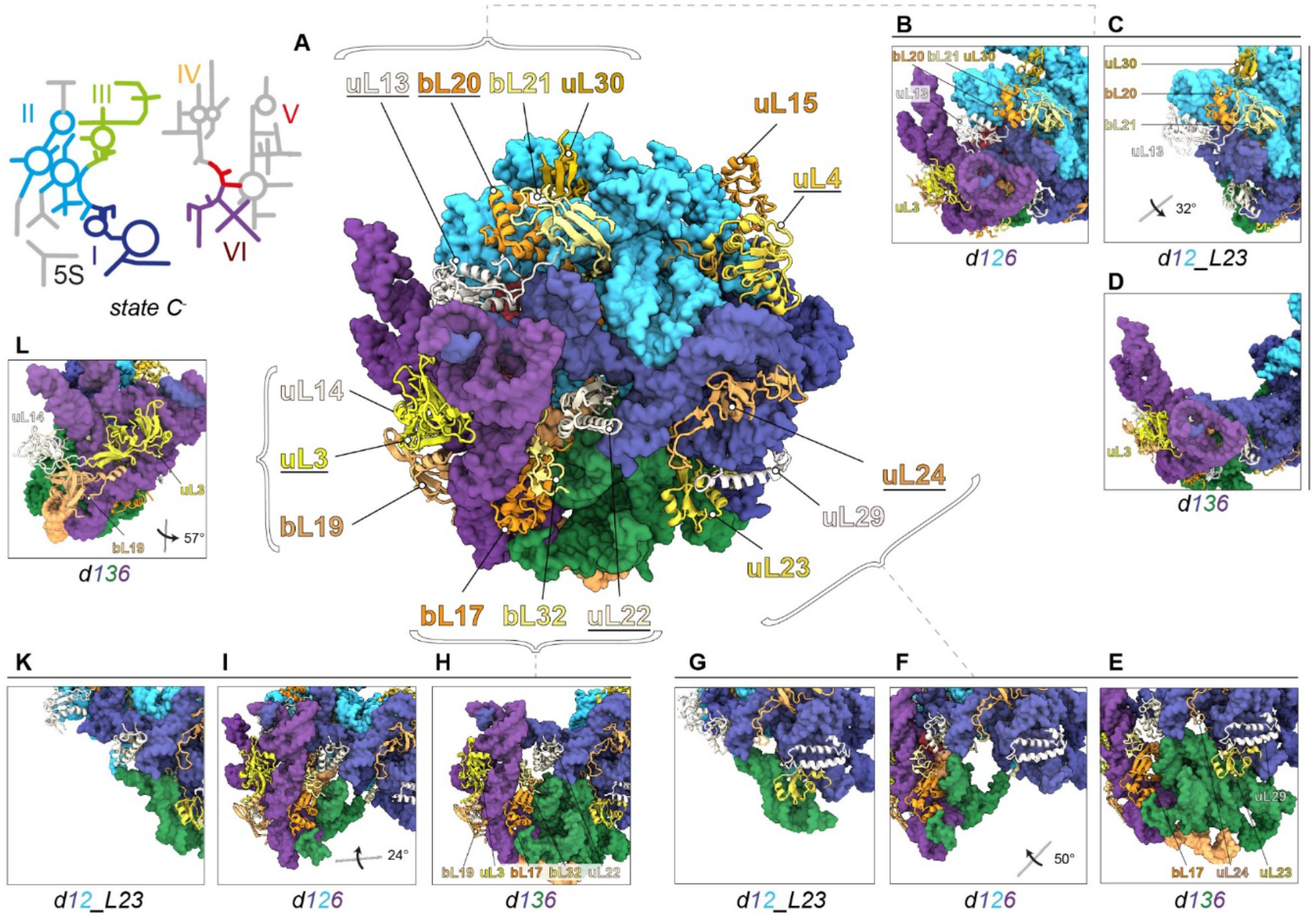
Clustering of L-proteins. **A)** 2D rRNA map and atomic model of state *C*^*-*^ in back view. L-proteins appear as cartoons (early essential proteins and uL3 are underlined), rRNA as surface model (lowpass-filtered to 5 Å resolution). Presence and absence of bL21, bL20 and uL30 in states *d126* **(B)**, *d12* **(C)**, and *d136* **(D)**, respectively. Presence and absence of uL23 in states *d136* **(E)**, *d126* **(F)**, and *d12* **(G)**, respectively. Presence and absence of bL17, bL32 and uL3 and bL19 in states *d136* **(H)**, *d126* **(I)**, and *d12* **(K)**, respectively. Positions of uL14, uL3 and bL19 in state *d136* **(L)**. Viewing angles in C), F), I) and L) are shown relative to the full model in A).

**Fig. 3:**
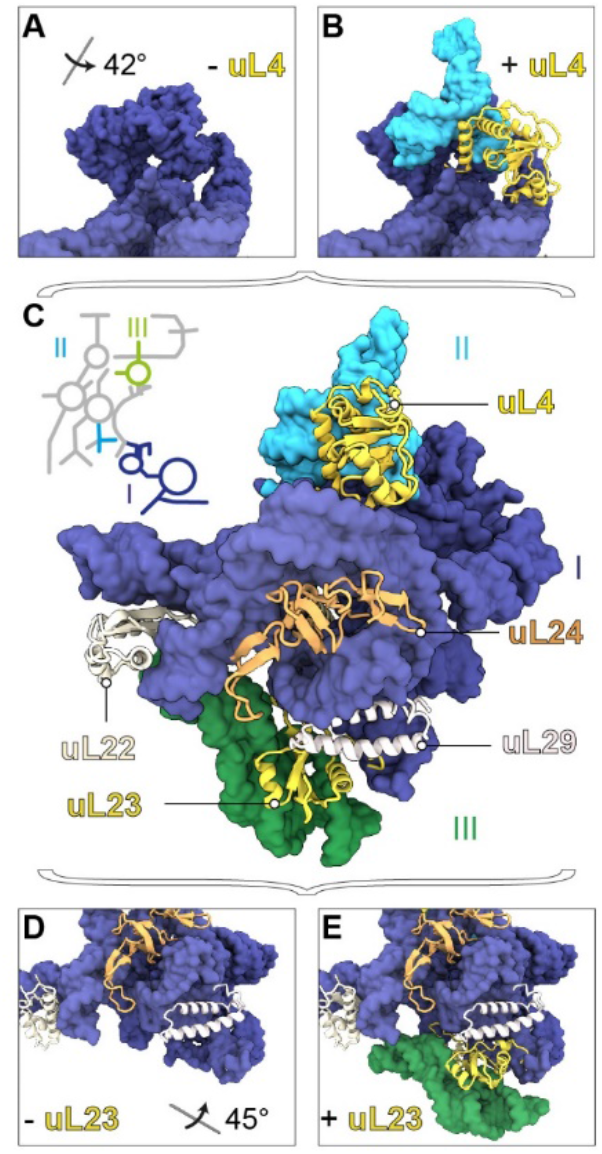
Proteins with seeding function. PDB models of early states with rRNA in surface representation and L-proteins as cartoons. Top segments of states *d1* **(A)** and *d1_L4* **(B)**, full model of state *d1_L4/L23* **(C)**, and bottom segments of states *d1* **(D)** and *d1_L4/L23* **(E)** color-coded according to the 23S rRNA 2D map. All viewing angles relative to (C).

Another L-protein that affects the formation of domain III is bL17, whose presence mediates stable folding of H47-49 **(Fig. 2H-K)**. Based on these observations, we conclude that some L-proteins, occupying surface exposed regions on a pre-50S particle, are capable of folding or, stabilizing rRNA elements of adjacent domains. While it is well-known that L-proteins are frequently located at rRNA domain interfaces ^6,7^, and that r-proteins have the capability to remodel rRNA in a cooperative fashion ^3^ this dedicated function of uL4, uL23, bL17 and other L-proteins in ribosome assembly was not apparent **(Movie 3)**.

### Detailed molecular morphogenesis – Successive manifestation of L-proteins

Globular domains of L-proteins, like in uL24, contribute to the formation of the subunit’s crust. Some L-proteins such as uL4, uL3, bL32, and uL2, contain a globular domain (yellow) combined with extended tentacles or loops (red) that interestingly project towards the center of the subunit **(Fig. 4A, B)**. The globular domain of uL4 appears early after formation of domain I, while most of the loop forms with domain II, and the apical tip even later with domain V **(Fig. 4C)**. The globular domain of uL3 critically contributes to formation of domain VI **(Fig. 2H, I, L)**, and the extended loop forms later with domains II, IV and V **(Fig. 4D)**. In addition, bL32 occupies a critical position with its globular domain being part of domain VI **(Fig. 2A, H, I)** and the extension interacting with domains I, II and V, thereby connecting four rRNA domains **(Fig. 4E)**.

**Fig. 4:**
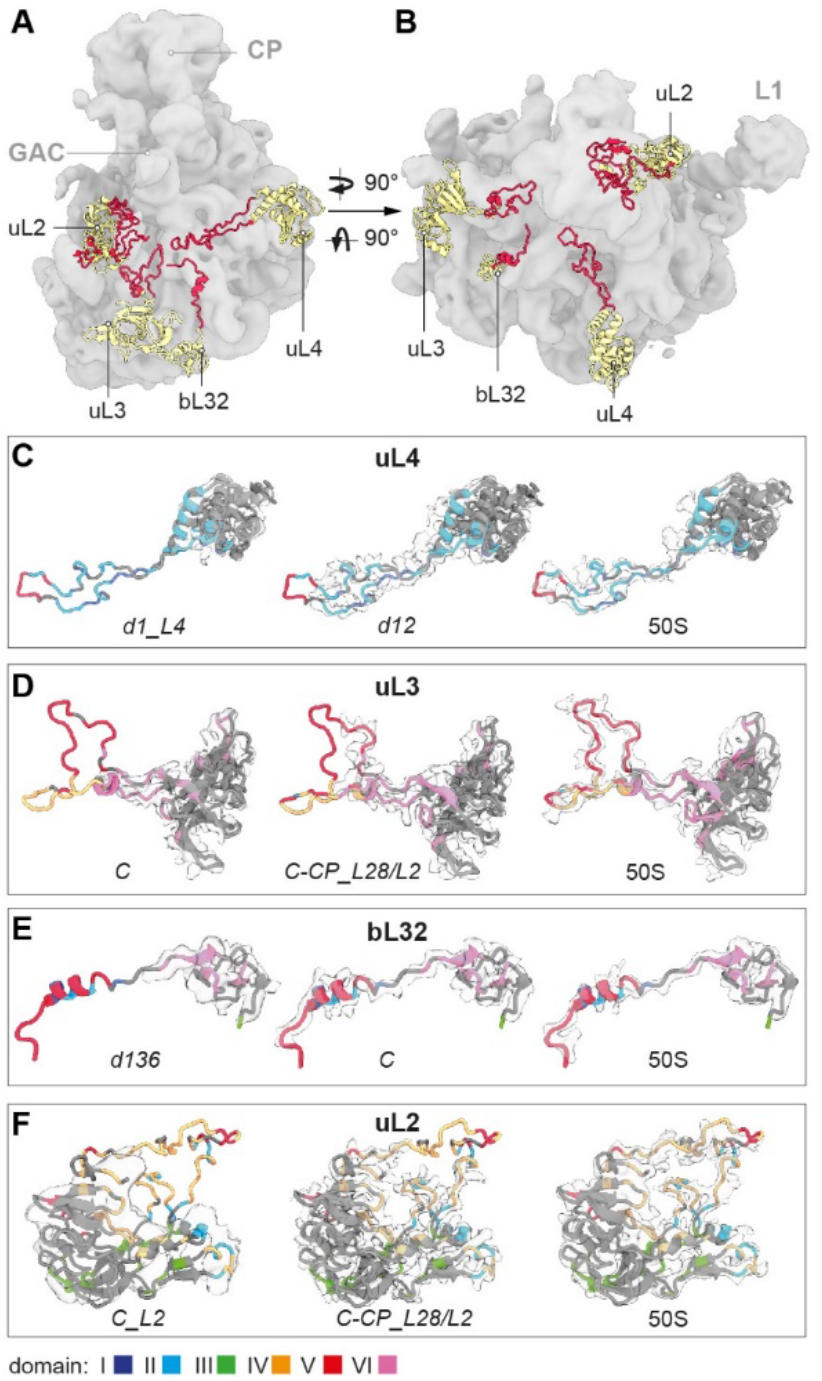
L-proteins with extensions. **A)** Side view, **B)** top view of a lowpass-filtered *C-CP-H68* cryo-EM map (gray) with models of selected L-proteins exhibiting globular domains (yellow) and extensions (red). CP, central protuberance; GAC, GTPase associated center; L1, L1 stalk. **C-F)** Cryo-EM maps (light gray) of selected L-proteins, derived from the indicated states, and corresponding atomic models (EMD-22614). Protein models are colored according to their rRNA domain contacts; gray color indicates self-interaction, or surface exposure (Material and Methods).

Interestingly, the globular part of uL2 is present on domain III only after core formation in state *C_L2* **(Extended Data Fig. 4M)** and binding can occur later in state *C-CP_L28/L2* **(Extended Data Fig. 4P and Extended Data Fig. 8A, D-G)**. Nevertheless, the extended loop starts forming earliest in state *C-CP_L28/L2* and interacts mainly with and stabilizes helices H65, H66 and H69 of domain IV. Only in a mature 50S subunit, density for the whole loop appears that in addition undergoes contacts with domain V **(Fig. 4F)**. Taken together, the flexible tentacles of some L-proteins form later than their globular domains and interact predominantly with nucleotides of the late forming rRNA domains IV and V, possibly by mechanisms involving dynamic sampling, as shown for proteins of the 30S subunit ^31,32^.

### Hierarchical arrangement of rRNA

Another principle we observe is the stabilization or folding of rRNA helices, induced by adjacent rRNA elements that are nearly maturely folded. Assembly appears to nucleate at the 5’-end with the formation of domain I to yield state *d1*. To rule out biased image classification by initial alignment, we re-analyzed our cryo-EM data and aligned all particle images to domain VI before proceeding with 3D classification **(Extended Data Fig. 6)**. However, this did not result in novel states that would support any nucleation point other than domain I. Thus, domain I indeed seems to appear first, while the remaining rRNA elements dock onto an already existing RNA surface.

The precursors *d12* **(Fig. 5A, B)** and *d136* **(Fig. 5C, D)** exhibit intrinsic flexibility along certain hinge regions (red dotted lines), while the individual rRNA domains remain relatively rigid. Nevertheless, the relative domain movements from the most open **(Fig. 5A, C)** to the close **(Fig. 5B, D)** conformation allow for the formation of inter-domain contacts between H11/ H37 **(Fig. 5B)** and H21/ H51 and H11/ H37 **(Fig. 5D)**. In some cases, formed helices remain flexible and are stabilized by tertiary contacts during subsequent assembly. For instance, both H21 and H22 exhibit solid density in state *C*^*-*^, but both helices reconfigure their shape and conformation and form tertiary contacts once reaching their mature positions **(Fig. 5E-G)**. Further rRNA-assisted rRNA stabilization or folding becomes apparent, when comparing assembly along the routes 1-2-6 and 1-3-6. While route 1-2-6 initially yields a core particle lacking density for domain IV helices H61-63 (state *C*^*-*^), maturation along route 1-3-6 culminates in a particle with these parts of domain IV stably folded (state *C* and better defined in state *C-CP_L2/L28*) **(Fig. 1D, and 5H-K)**. Hence, initial appearance of domain IV benefits from the presence of the maturely folded domain III. Specifically, rRNA contacts with the already formed domain III helices H56-58 seem to induce formation of domain IV H61 and H63 in state *d136*, and H61-63 in state *C* **(Fig. 2A-C and Extended Data Fig. 4K, L)**, thereby initiating the incremental formation of domain IV.

**Fig. 5:**
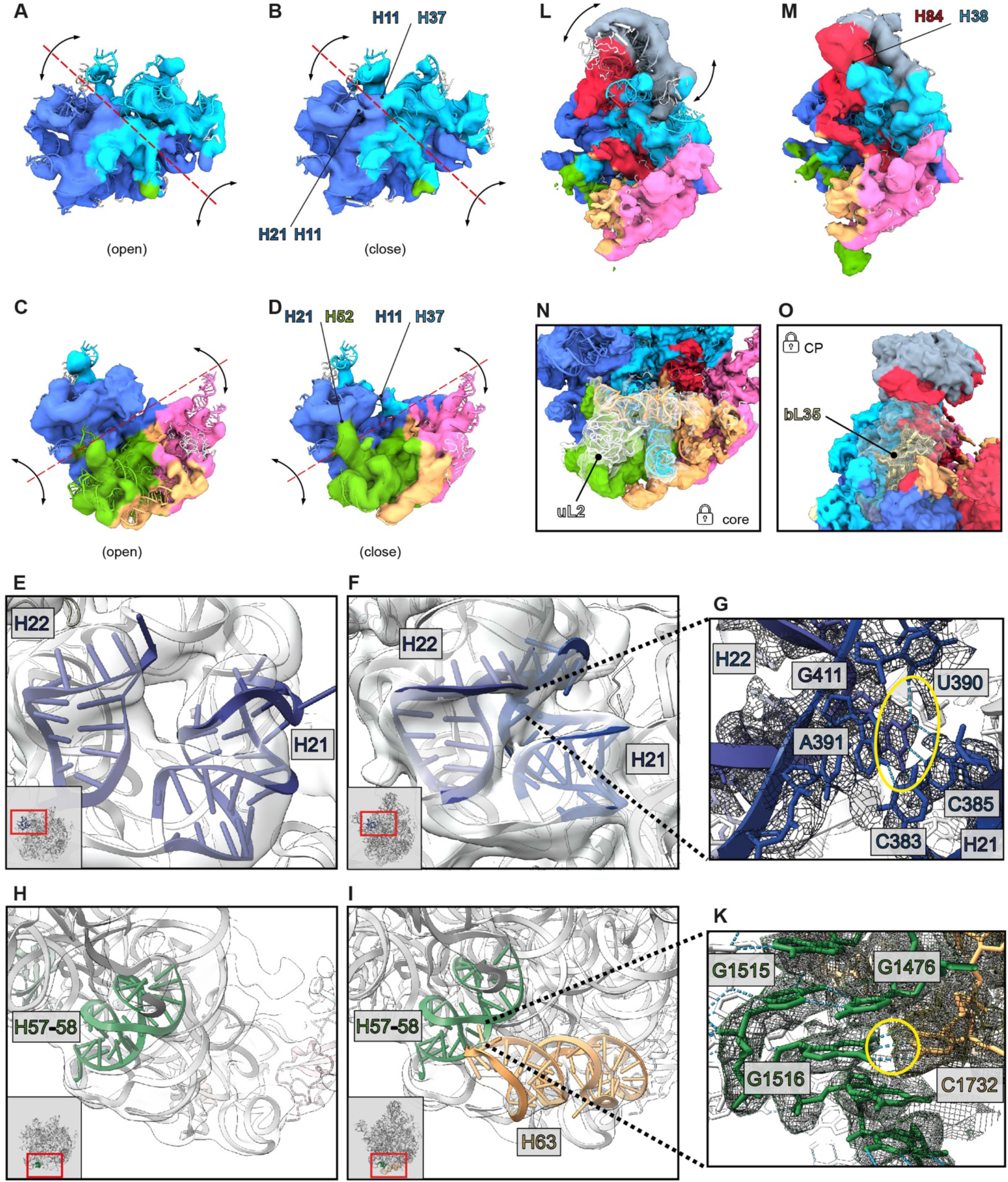
Tertiary contacts between rRNA helices. 3D variability maps of states *d12* **(A, B)**, d136 **(C, D)**, *d126-CP* **(E, F)**. First (left) and last (right) reconstructed frame of each state is shown. Maps were filtered to 8 Å and color-coded according to the six architectural domains of 23S rRNA. Red dotted lines and black arrows indicate axis and directions of observed global movements, respectively. Stabilizing contacts in closed conformations are indicated by black arrows. Interacting regions are labeled. Binding sites of L-proteins uL2 (left, state *C_L2*) and bL35 (right, state *C-CP-H68*) are shown **(G, H)**. Cryo-EM maps (transparent gray) and PDB models corresponding to rRNA helices H21 and H22 (blue), H57-58 (green) or H63 (orange), respectively. Local snapshots of state *C*^-^ **(I, M)** and state *C-CP_L2/L28* **(K, N)** contain thumbnails with red rectangles indicating the region of interest. **L, O)** close-ups of regions of interest in state *C-CP_L2/L28*. Yellow circles highlight H-bond interactions (light blue dotted lines) between C385-G411 **(L)** and G1516-C1732 **(O)**.

In addition, we noticed that the 5S RNP (ribonucleoprotein particle), consisting of 5S rRNA, uL5, uL18 and bL25, can dock on a pre-50S particle as soon as the central region of domain II has formed **(Extended Data Fig. 7A, B)**. This can occur both in early (*d12-CP, d126-CP*) and late assembly (*C-CP*) and goes along with formation of domain II helices H38 (A-site finger) and H42 (GAC), as well as domain V helices H80-H88 **(Extended Data Fig. 4D, F, N, and Extended Data Fig. 7C)**, representing additional examples for rRNA induced rRNA stabilization or folding. However, the CP remains mobile and upon rotational movement of the 5S rRNA towards the subunit interface, H84 and H38 stabilize each other **(Fig. 5 L&M)**.

### Late 50S assembly

Late 50S assembly, apart from core-stabilizing incorporation of uL2, involves stable binding of bL9 and bL28 in state *C-CP_L28*, which rigidifies the L1 stalk **(Extended Data Fig. 4O and Extended Data Fig. 8D-I)**. Next, H68 can form upon integration of bL33 and bL35 **(Extended Data Fig. 4Q and Extended Data Fig. 8K-L)**. In addition, the fixation of H68 seems to involve contacts with uL1. Furthermore, the L1-stalk associated H75 and CP associated H88 provide RNA contacts and stabilize H68. Accordingly, late step 1 assembly terminates with the “48S” particle *C-CP-H68*, sharing most structural features with a mature 50S subunit, but lacking a properly arranged FC.

## Discussion

Here we show, using the 50S *in vitro* reconstitution assay in combination with cryo-EM and multi-particle refinement that already after 3 min of reaction as much as 16 structurally distinct precursors have formed. Hence, the structures we report provide unprecedented insights into the detailed progression of 50S *in vitro* assembly, addressing long-standing questions and revealing fundamental principles of large subunit assembly.

One important question related to *in vitro* ribosome assembly assays is, how subunit assembly can take place, despite the presence of full 16S or 23S rRNA? In principle, structure formation could initiate at multiple, possibly competing sites. While we cannot rule out this scenario (based on our qMS data), we did not obtain structural evidence for any nucleation point other than domain I **(Extended Data Fig. 6)**. Instead, our data indicate that the assembly of the 50S subunit does not proceed arbitrarily but has its morphogenetic origin in domain I.

Based on their intrinsic affinities, uL22, uL24 and uL29 interact with their RNA binding sites in domain I and nucleate 50S assembly. Consequently, binding sites for uL4 and bL23 are formed. Next, binding of uL4 and bL23 initiates formation of domains II and III, respectively, and further proteins join, exemplifying the inherent cooperativity of ribosome assembly. Hence, we visualize Nomura’s paradigm, that all the information for ribosome assembly is contained in the structure (i. e. the chemical affinities) of the participating components ^33^. As a practical consequence, atomic models derived from the assembly intermediates will be useful for the design of single molecule approaches and pharmacological agents interfering with critical transitions along the 50S assembly pathway.

After formation of domain I, assembly continues uL4 dependent with *d1_L4* and branches either to route 1-2-6 or 1-3-6 **(Fig. 6)**. The presence of domain II apparently facilitates docking of the CP-forming 5S RNP and as a consequence, route 1-2-6 involves the possibility of CP formation either on *d12*, or *d126* resulting in *d12-CP*, or *d126-CP*, respectively. However, completion of the CP can also occur after formation of state *C* during incremental late assembly **(Extended Data Fig. 5)**. Route 1-3-6 yields state *C* due to presence of domain III, which is seeded by uL23, and in turn facilitates incorporation of domain VI independently of domain II **(Fig. 6, and Extended Data Fig. 5)**. Furthermore, additional routes are conceivable. Precursor *d12_L23* could derive from both *d12* and *d1_L4/L23* and constitute an intermediate that matures to state *C* by completion of domain III and simultaneous or subsequent formation of domain VI. Proteins that contribute to the formation of these domains are bL17, bL32 and uL3 **(Fig. 6)**.

**Fig. 6:**
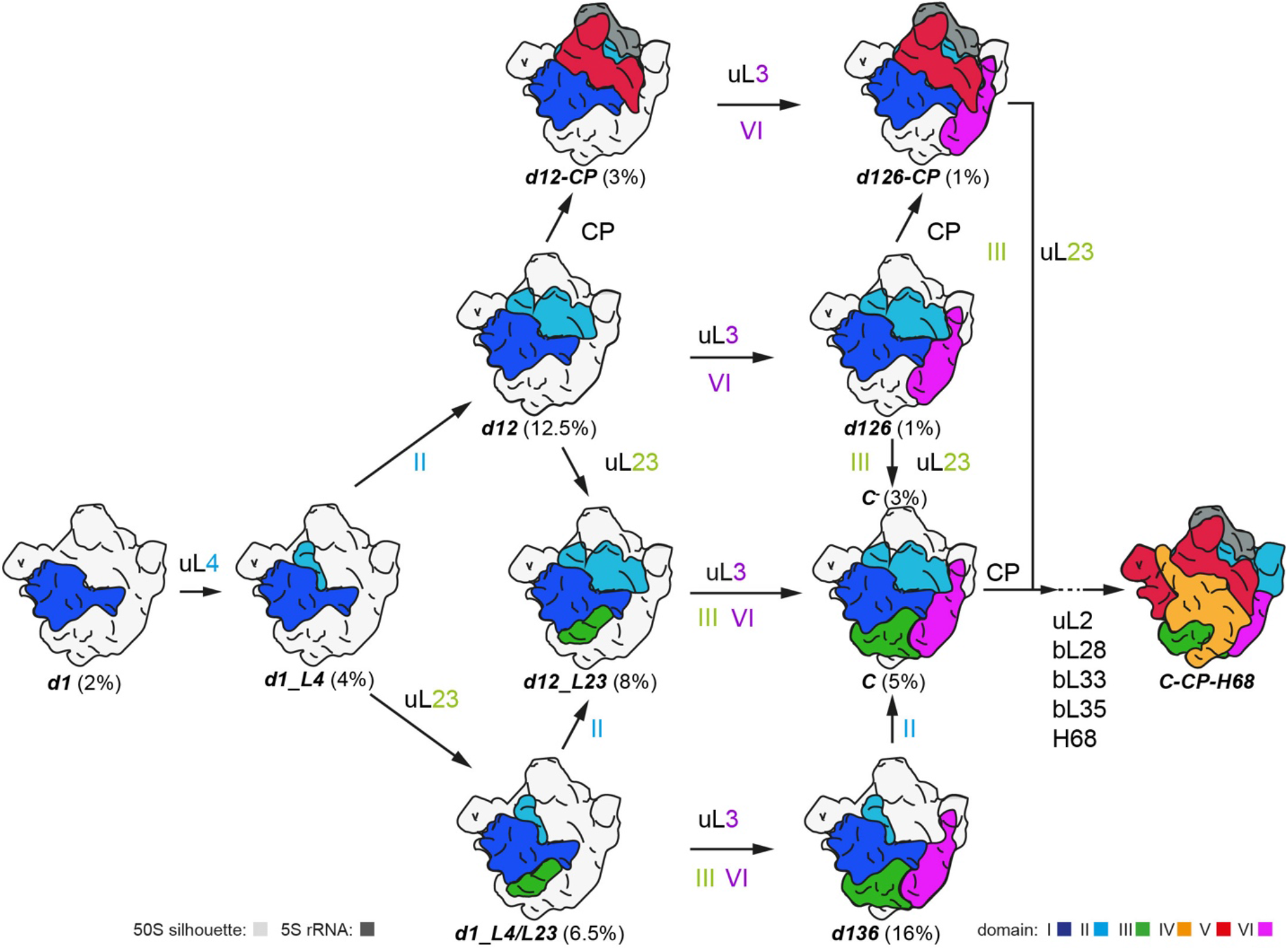
Model of early 50S assembly. Models of individual states color-coded as indicated. Numbers in % represent their relative abundance in the sorting scheme. Gray silhouette illustrates density of the mature 50S subunit. The labels uL4, uL23, uL3 and the numbers II, III and VI indicate when cryo-EM densities for parts of these proteins or entire 23S rRNA domains, respectively, become detectable. CP, central protuberance; H68, helix 68.

Accordingly, early 50S assembly is characterized by hierarchical arrangements of rRNA, resulting in branches, but on the other hand generates cooperativity and ultimately culminates in convergence along the assembly pathway. Due to the interwoven architecture and the ring-shaped arrangement of domains I, II, III and VI, domain VI can be incorporated via routes 1-2-6 or 1-3-6. While uL24 promotes formation of the 5’-domain I, the late assembling uL3 plays a critical role in the structural organization of the 3’end-containing domain VI. As we do not observe a state consisting only of domains I and VI, their contact areas may be too small to drive stable incorporation of domain VI in the absence of either domain II or III. This suggests that domain I together with domain VI cradle the missing domain in *d126*, or *d136*, to promote appearance of domain III or II, respectively, to conclude core formation and early assembly **(Fig. 6)**. Thus, our analysis indicates that coordinated structural formation of the two terminal domains of 23S rRNA is a decisive event that drives further assembly.

Structural data regarding earliest stages of large subunit assembly are lacking from both the prokaryotic and the eukaryotic domain. Interestingly, a structural analysis of nucleolar pre-60S assembly captured a state, consisting of 28S rRNA domains I and II (similar to our state *d12*), that was thought to proceed linearly to a 60S subunit ^27^. Surprisingly, a second study found evidence for parallel assembly pathways ^28^. While we find similar states, confirming the evolutionary conservation of the process, our analysis starts with a particle whose assembly has just initiated (*d1*). In addition, we can take advantage of the multitude of states we were able to obtain after thorough sorting and trace how decisive steps are made towards a mature 50S subunit, involving multiple parallel routes of assembly.

Taken together, the combination of time-limited *in vitro* 50S assembly and cryo-EM allowed us to obtain unprecedented insights into the early assembly phase of the bacterial large ribosomal subunit. However, to obtain a more detailed understanding of the assembly process, fluorescence-based real time experiments and approaches clarifying the mechanistic role of ribosome assembly factors will be required.

## Materials and Methods

### 50S *in vitro* reconstitution

The assay was performed according to ^34^ with the modifications described in ^14^. In step 1 of the 50S reconstitution, seven samples were prepared in individual tubes corresponding to 7 time points. In each of the tubes, 6.25 equivalent units of TP50 (total protein of the 50S subunit) and 6.25 A^260^ rRNA (23S rRNA + 5S rRNA) were mixed on ice and incubated at 44 °C using a thermo-block (Eppendorf, Thermomixer comfort) in Rec4 buffer (20 mM HEPES/KOH pH7.6, 400 mM NH^4^Ac, 4 mM Mg(Ac)^2^, 0.2mM EDTA, 5mM 2-mercaptoethanol). A first aliquot was taken instantaneously (0 min) and further samples were taken after 2, 3, 5, 10, 20 and 30 min of incubation. 36 pmol of material from the individual time points were adjusted to 20 mM Mg(Ac)^2^ concentration and incubated for 90 min at 50 °C.

### Analytical sucrose density gradient ultra-centrifugation

18 pmol material from the individual time course were subjected to analytical sucrose density gradient (10%–30%) ultra-centrifugation (SW40 rotor, 26,000 rpm for 17 h) in Tico buffer (20 mM HEPES/KOH pH7.6 on ice, 30 mM KAc, 6 mM Mg(Ac)^2^, 4 mM 2-mercaptoethanol).

### *In vitro* translation assay (poly(U)-dependent poly(Phe) assay)

6 pmol isolated native 50S subunits (50S), reconstituted 50S subunits (50S rec) from each time point of step 1 that had been subjected to a full step 2 incubation, were mixed with 12 pmol 30S subunits and incubated in assay buffer (20 mM HEPES/KOH pH7.6, 150 mM NH_4_Ac, 6 mM Mg(Ac)_2_, 0.05 mM EDTA, 4 mM 2-mercaptoethanol, 0.05 mM spermine, 2 mM spermidine, 3 mM ATP, 1.5 mM GTP, 5 mM acetyl-phosphate, 83 µM [^14^C] Phe (22 dpm/pmol) (Hartmann analytic GmbH), 0.83 mg/ml poly(U), 0.34 mg/ml tRNA^bulk^ and S100 extract) for 60 min at 37 °C. Subsequently, 30 µl BSA (1% stock solution) and 2 ml TCA (10%) were added. Samples were briefly vortexed, incubated for 15 min at 90 °C and 5 min on ice. Precipitated proteins were immobilized on glass filters, washed two times with 5% TCA and once with ether/ethanol (1:1). The filters were incubated in scintillation liquid (Roth, Rotiszint eco plus) and the amount of incorporated ^14^C Phe was determined using a scintillation counter (Wallac 1409 liquid scintillation counter). The assay was performed in duplicates and mean values were calculated.

### Quantitative mass spectrometry analysis

108 pmol of the individual samples from the step 1 time course reaction were adjusted to total volume 180 µl and loaded on 900 µl sucrose cushion (20% in Tico buffer). Ultra-centrifugation was performed (TLA-110 rotor, 65,000 rpm for 2.5 h). After removing the supernatant, the pellet was briefly washed and dissolved in Tico buffer. 36 pmol of each sample were analyzed by quantitative mass spectrometry according to the workflow described in ^12^. For protein comparisons per sample, log transformed iBAQ intensities were normalized to uL24, which is an early assembly protein of the large bacterial ribosomal subunit. Fold changes to the reference purified mature 50S was calculated from normalized iBAQ values. The ribosomal stoichiometry of individual L-proteins in each sample was calculated as the percentual fraction of the value of corresponding L-proteins in mature 50S. Therefore, 100% stoichiometry represents a protein that is as abundant as its counterpart in the mature 50S subunit. Experiments were performed in duplicates and data were processed with Heatmap Illustrator 1.0 (CUCKOO Workgroup) and Origin8 (OriginLab Corporation).

### Sample preparation for cryo-EM

12 pmol of each sample purified via sucrose cushion ultra-centrifugation, as described above, were diluted twofold in adaptation buffer (20 mM HEPES/KOH pH7.6, 4 mM Mg(Ac)_2_, 0.2 mM EDTA, 5 mM 2-mercaptoethanol) to achieve a final concentration of 288 pmol/ml and spotted on glow-discharged holey carbon grids (R1.2/1.3 copper 400 mesh holey carbon, without additional carbon film on top (carbon-free)), (Quantifoil Micro Tools GmbH) and cryo plunged in liquid ethane after blotting using a Vitrobot device (MK4).

### Cryo-electron microscopy and data processing

Data were collected on a FEI TecnaiG2 Polara equipped with a Gatan K2 Summit detector operated in super-resolution mode at 300 kV at a calibrated pixel size of 0.625 Å. Movies were acquired for 5 sec applying a total electron dose of 25 e^-^/Å^2^.

### Data processing

Movies were aligned and dose-weighted using MotionCor2 ^35^. Defocus values were estimated using Gctf ^36^. Particles were picked using templates in Gautomatch (developed by K. Zhang). Templates were generated in SPIDER ^37^. Therefore, density maps were generated from atomic models of state 1 (PDB: 6GC7) and state 5 (6GBZ), subsequently low-pass filtered to 20 Å, followed by projection into 84 equally distributed orientations, respectively. Orientation images were averaged into four projections using Xmipp3 2D classification ^38^ classification for each template, combined into one stack and then used for particle picking. Particle images were extracted and normalized, using Relion 3.0 ^39^ with a box size of 600 and Fourier cropped to 150 for sorting. If not stated otherwise, Cryosparc v3.1 ^40^ was used for the identification and refinement of final classes. Initial sorting was achieved using ab-initio classification, followed by two rounds of heterogeneous refinement to recover ribosomal particles. Remarkably, recovered ribosomal particle classes exhibited density for domain 1 and 2 of the 23S rRNA in the first and density for domain 1 only in the second heterogeneous refinement. Particles were then pooled and aligned to a ribosomal consensus map, generated in the initial ab-initio classification. Remaining particles were sorted out using reference-free 2D classification. Identification of main classes were performed using 3D classification without alignment in Relion 3.1. Best classification results were achieved, when particles were first locally refined to domain 1 of the 23S rRNA. Subsequently, classes were validated, reassigned and further sorted using identified classes and heterogeneous refinement. An additional 3D ice template was generated by ab-initio reconstruction using identified classes in the 2D classification. Remaining classes were identified using hierarchical 3D variability clustering. When necessary, additional rounds of multi-class ab-initio reconstruction (class similarity = 0) or heterogeneous refinement were carried out to identify shiny particles. The most mature precursor, *C-CP-*H68, still showed marked fragmented density for uL16, uL35 and H68, indicating remaining structural heterogeneity. However, due to low particle numbers, it was not sorted further. After completion of particle classifications, sub-classes were refined using non-uniform refinement at a pixel size of 1.25 Å.

### Model building

Atomic models of previously identified 50S assembly intermediates (PDB: 6GC0, 6GC4, 6GC7, 6GC8) were used for modeling. Initially, models were rigid body docked in ChimeraX ^41^, followed by rigid body fitting of individual L-proteins. RNA regions and L-proteins were removed, when densities were missing or strongly fragmented. The adjustment of structured elements was performed by iterative model building and real-space refinement into the EM-density using Coot 0.9.6 ^42^, Phenix 1.19 ^43^ and ERRASER ^44^ considering secondary structure restraints. For the analysis of L-protein mediated sampling (Fig. 3), full-length models of respective L-proteins were fitted into EM maps of indicated states. A 50S Cryo-EM map (EMD-222614) was used for comparison ^45^. Models were colored according to any rRNA contacts within 4 Å distance and the corresponding 23S architectural domain. Self-contacts and surface exposure was colored in gray. Local Cryo-EM maps are shown at 3Å distance to atomic models using the ChimeraX zone tool. For better visualization, noise and small fragmented regions were excluded by using the surface dust tool.

## Acknowledgements

We would like to thank Anett Unbehaun, Timo Fluegel and Matthew Kraushar for intensive scientific discussions, excellent suggestions, and text editing.

## Author Contributions

B.Q. performed all biochemical experiments and prepared sample for cryo-EM and qMS; S.M.L. processed, analyzed, and visualized structural data and built atomic models. A.B. and P.S. evaluated and improved atomic models; C.H.V.-V. and M.S. performed qMS and analyzed the data. J.B. and T.M. participated in the grid preparation, operation of the microscopes, data acquisition and processing; C.M.T.S. and R.N. supervised the study. R. N. drafted the manuscript. All authors contributed to writing of the manuscript.

## Data and materials availability

All data needed to evaluate the conclusions in the paper are present in the paper and/or the Supplementary Materials.

## Supplementary Materials for

**Extended Data Fig. 1:**
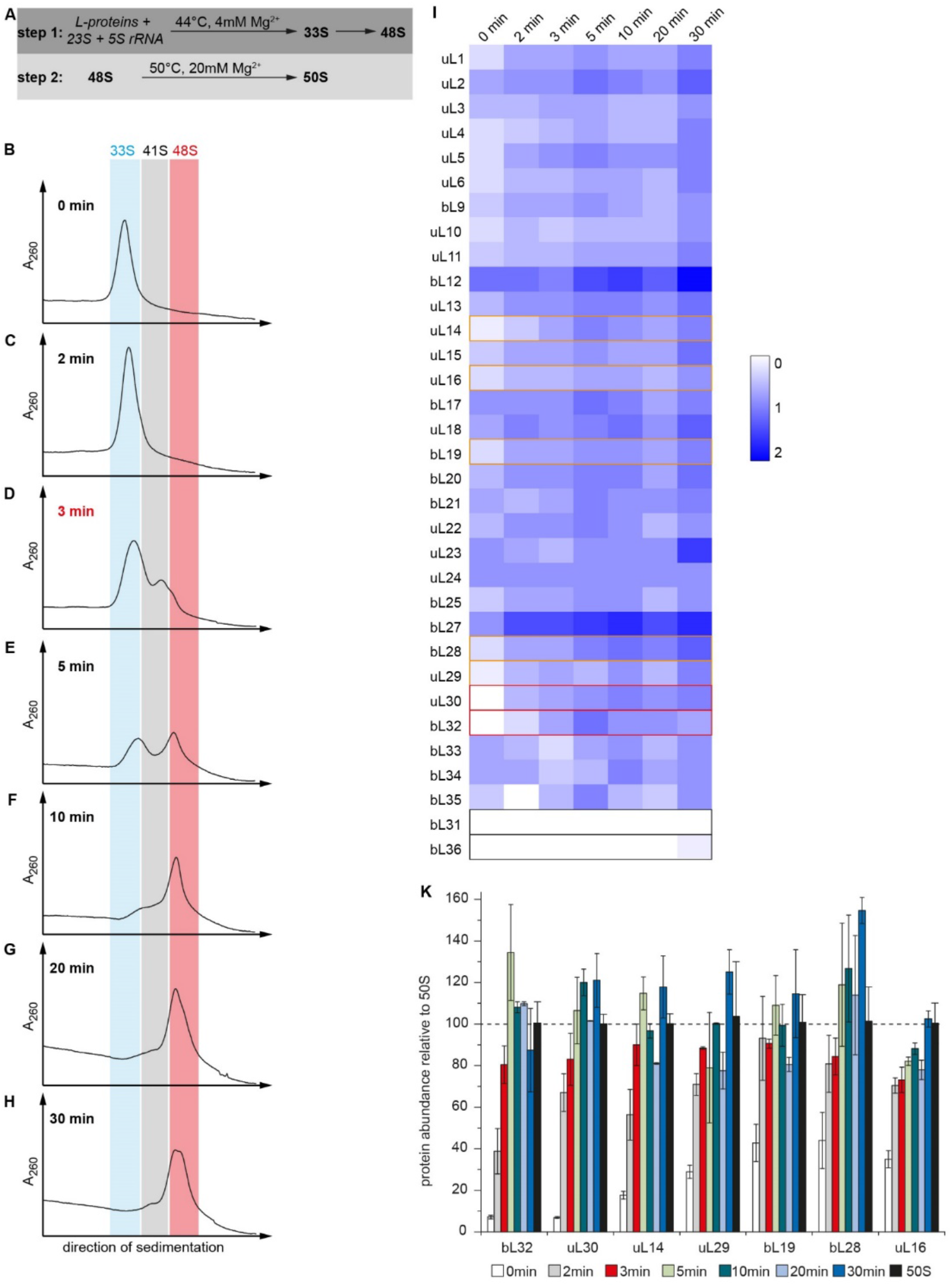
50S *in vitro* reconstitution assay and subsequent analyses. **A)** Two step *in vitro* reconstitution assay of the 50S subunit. In step 1, 23S and 5S rRNA together with L-proteins incubate at 44°C for up to 30 min and are spontaneously converted into a 33S particle, which ultimately matures to 48S particles. During step 2, 48S particles (material in the red zone (b-h)) are quantitatively transformed into 50S like particles after 90 min at 50°C. **B-H)** step 1 time course reaction. Samples incubated under step 1 conditions as indicated were subjected to sucrose density gradient ultracentrifugation. **I, K)** quantitative mass spectrometry analysis of material incubated under step 1 conditions, subjected to sucrose cushion purification and LC-MS. Relative abundance of all L-proteins plotted as heat map **(I)** and selected L-proteins plotted as bar chart **(K)**. Experiments were performed in technical duplicates and results represent mean values with error bars depicting standard deviation.

**Extended Data Fig. 2:**
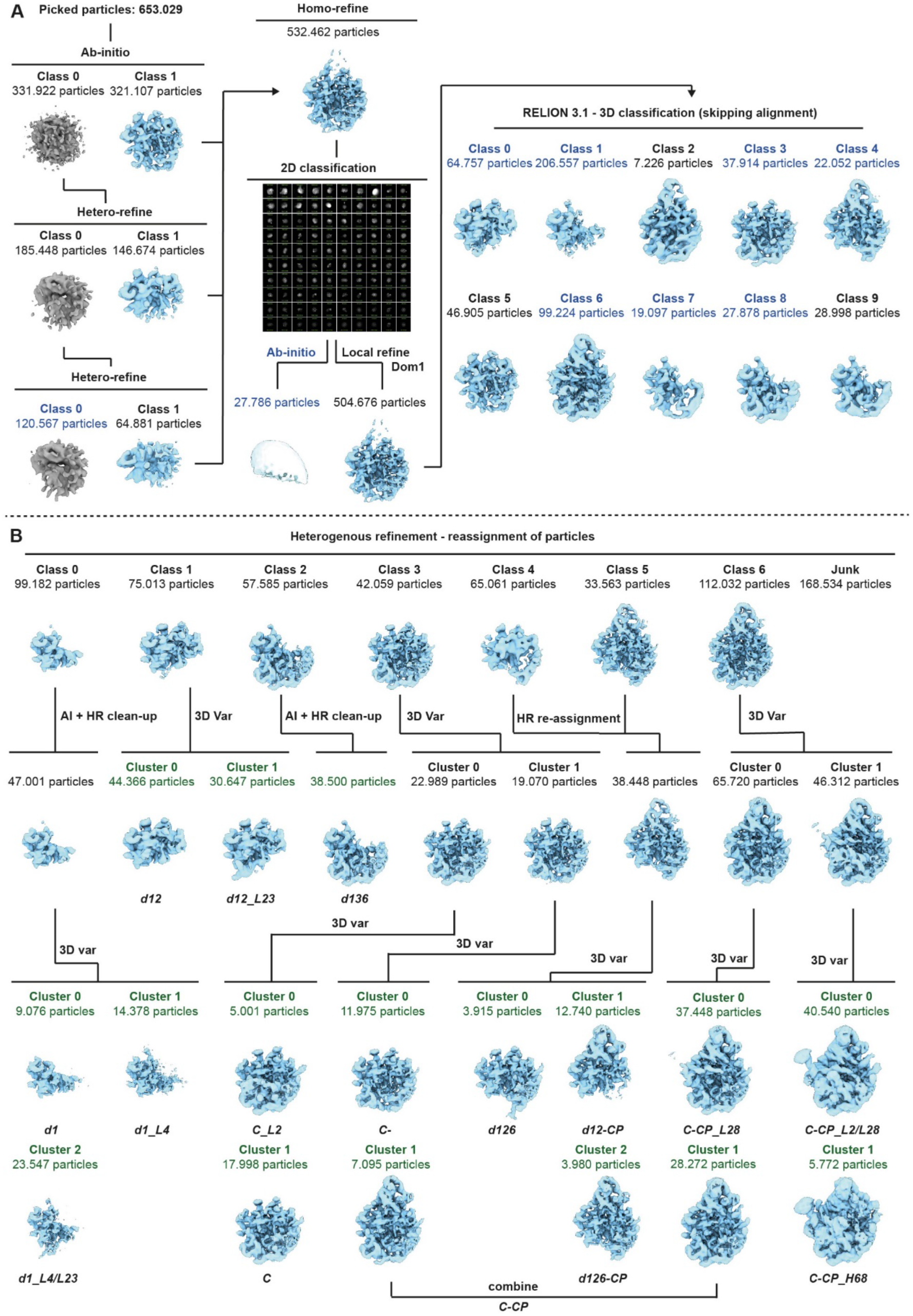
Sorting scheme. **A)** Sorting of particles and identification of initial classes. Extracted particles were subjected to an ab-initio reconstruction, followed by two rounds of heterogeneous refinement to recover ribosomal particles. Particles were refined to a consensus map and remaining ice particles were sorted out using 2D classification. Particles were aligned to the 23S rRNA domain 1 region using local refinement and sorted using 3D classification skipping alignment in Relion 3.1. Templates used for particle re-assignment are labeled in blue. An ice template was generated using an ab-initio reconstruction from selected 2D classes. **B)** Re-assignment of initial classes, cleaning and identification of final classes (green). Particles were re-assigned to identify initial classes. Further sub-classes were identified using 3D variability clustering. For *d1* and *d136* classes, assigned particles were cleaned using ab-initio reconstruction followed by heterogeneous refinement to identify most defined subsets. Sorting was performed at a pixel size of 3.75 Å. Final classes were refined at a pixel size of 1.25 Å (Extended Data Fig.3).

**Extended Data Fig. 3:**
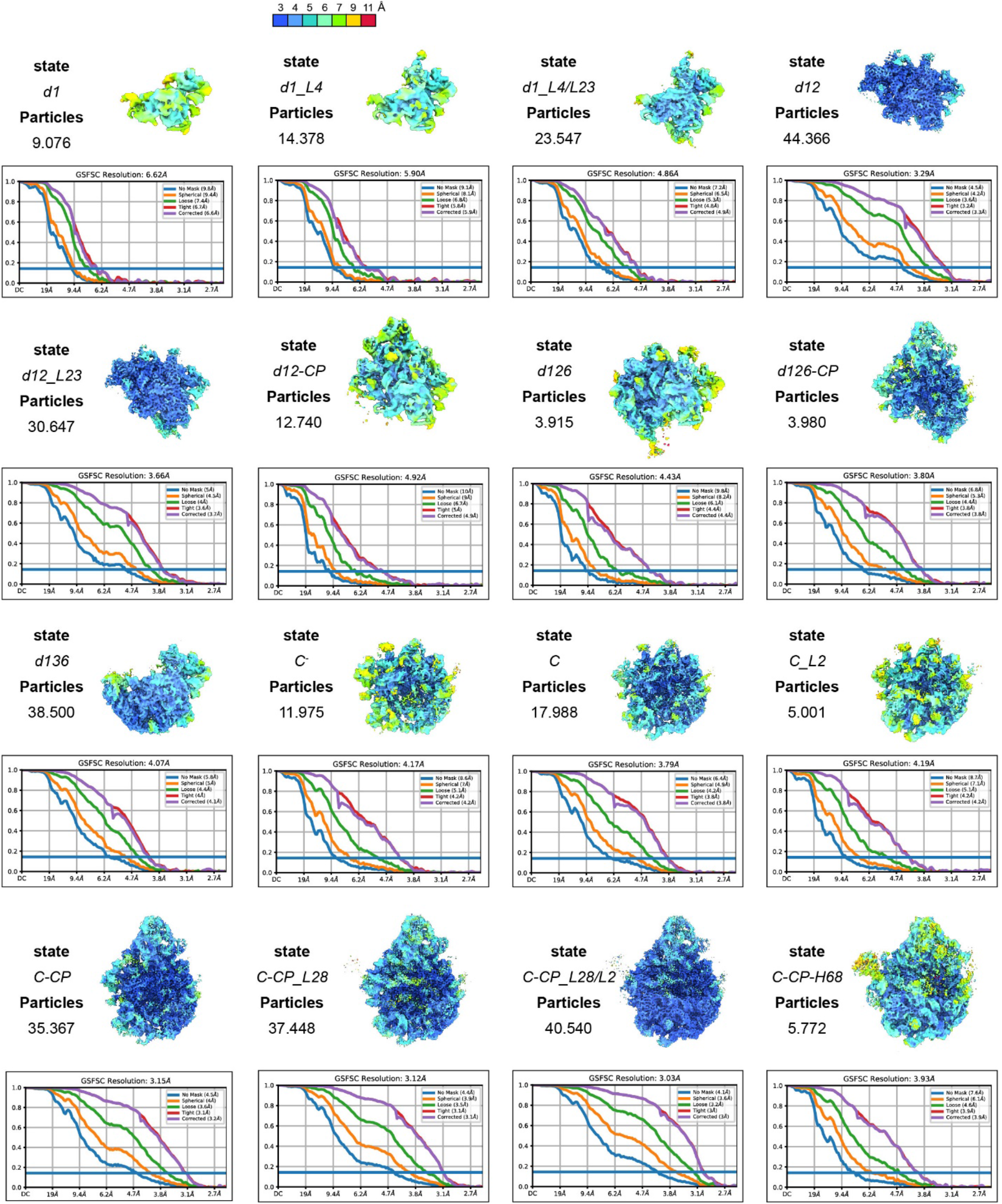
Final Cryo-EM maps and validation. Local resolution maps, total number of particles and gold-standard Fourier shell correlation (GSFSC) plots are shown for each state. States *d1, d1_L4, d1_L4/L23, d12, d12_L23, d12-CP, d126* and *d136* shown in back view, states *d126-CP, C*^*-*^, *C, C_L2, C-CP, C-CP_L28, C-CP_L28/L2* and *C-CP-H68* shown in crown view. Local resolution is shown using color keys from 3 Å (blue) to 11 Å (red). Overall resolutions were calculated from half-maps using the GSFC criteria cutoff of 0.143.

**Extended Data Fig. 4:**
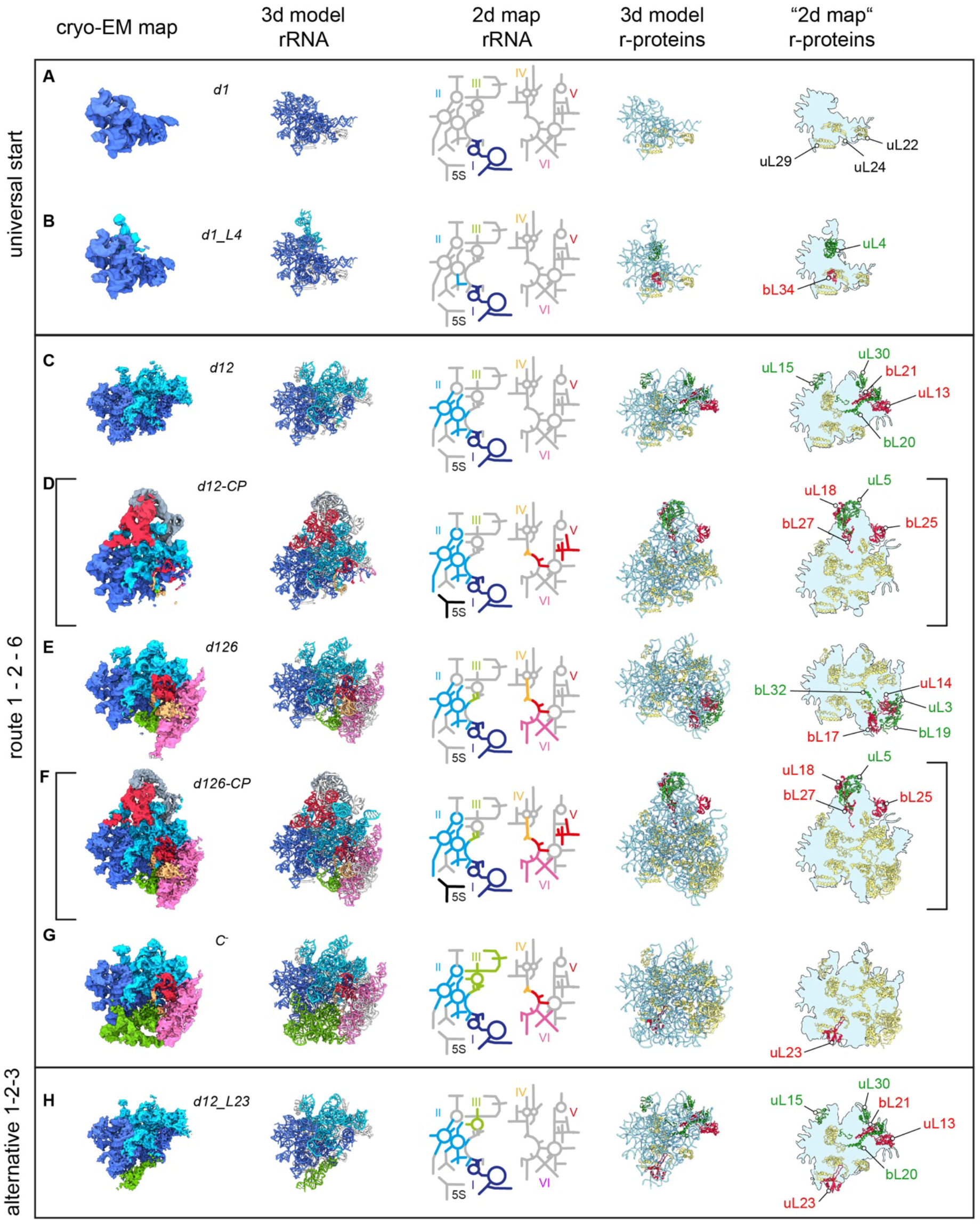
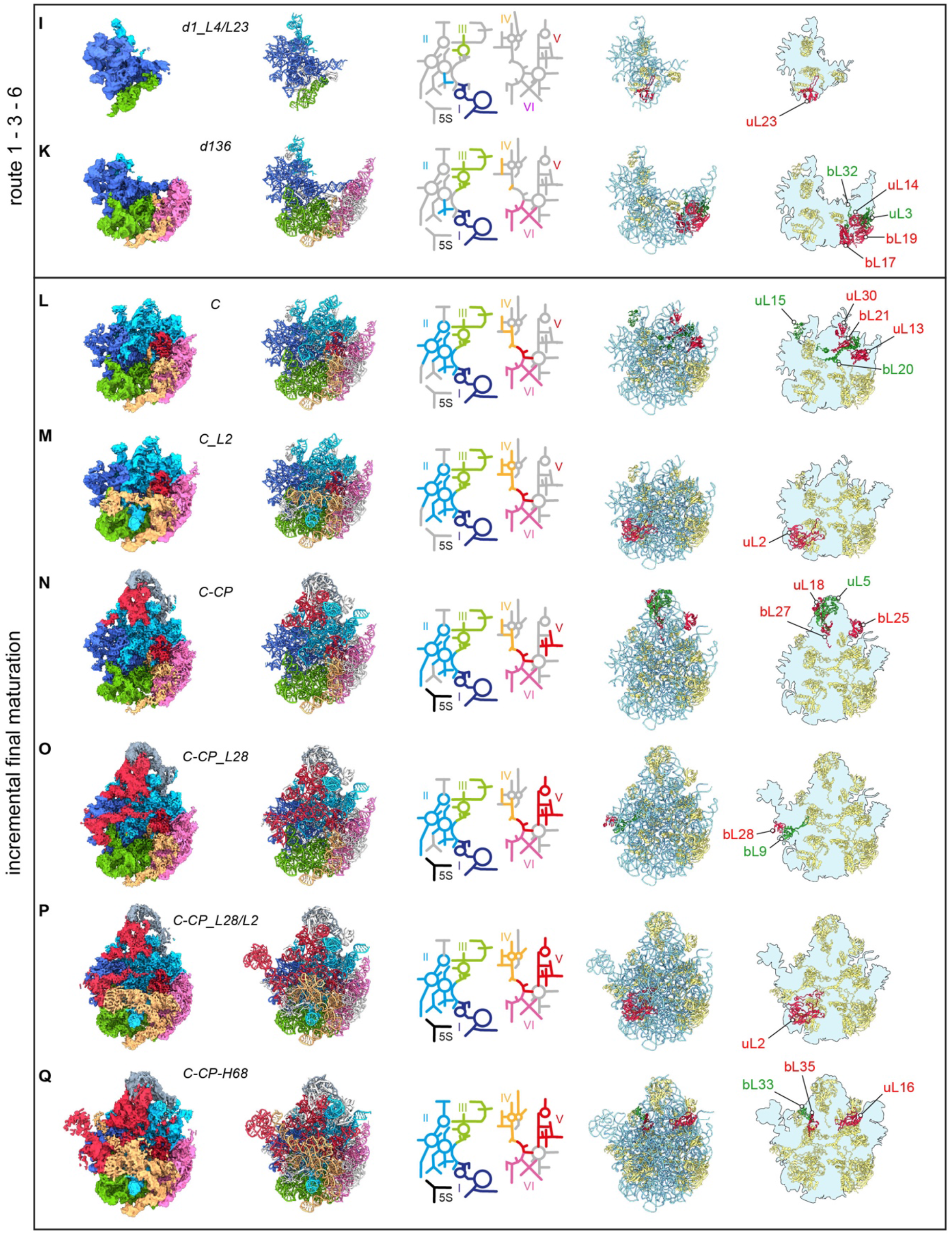
Gallery of pre-50S states. **A-Q)** States were lined up by their degree of structural completeness and their routes of maturation. Cryo-EM maps with rRNA and L-proteins color-coded according to the six architectural domains of 23S rRNA they are part of. 3D PDB models, and 2D maps with the six rRNA domains color-coded. Colored regions indicate stably formed segments, for which cryo-EM densities were obtained. 3D PDB models, and “2D maps” with rRNA in pale blue and L-proteins in gold. Proteins for which density appears compared to the previous state are highlighted in red or green. **D and F)** Early assembly intermediates with cryo-EM density for the CP, *d12-CP* and *d126-CP*, respectively.

**Extended Data Fig. 5:**
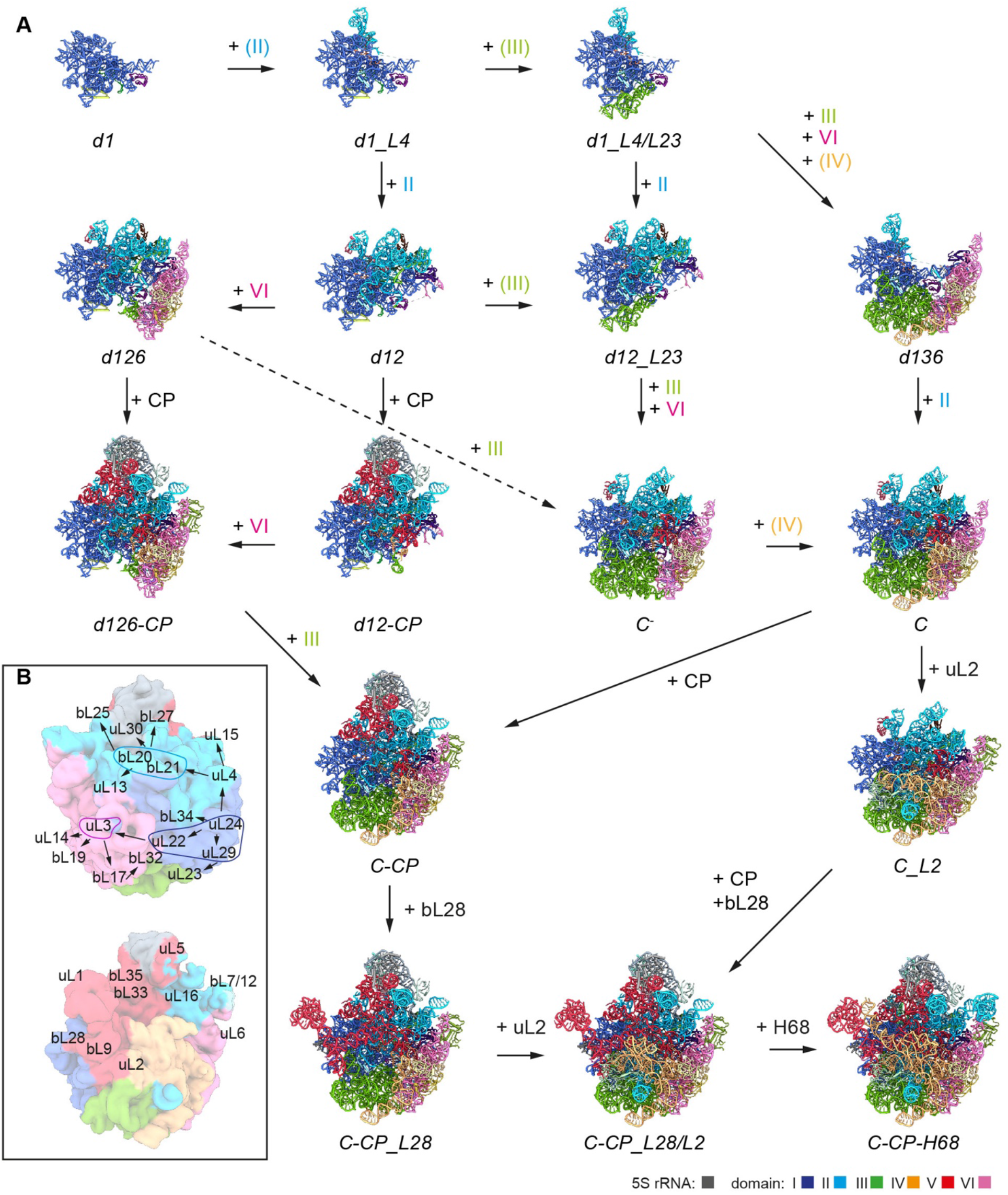
Interconversion of pre-50S states. **A)** PDB models of the individual states, with rRNA color-coded as specified. Sequences of appearing features (L-proteins or rRNA elements) are indicated. **B)** Possible order of L-protein binding, starting with uL24. The rRNA domains in brackets indicate partial formation of these domains.

**Extended Data Fig. 6:**
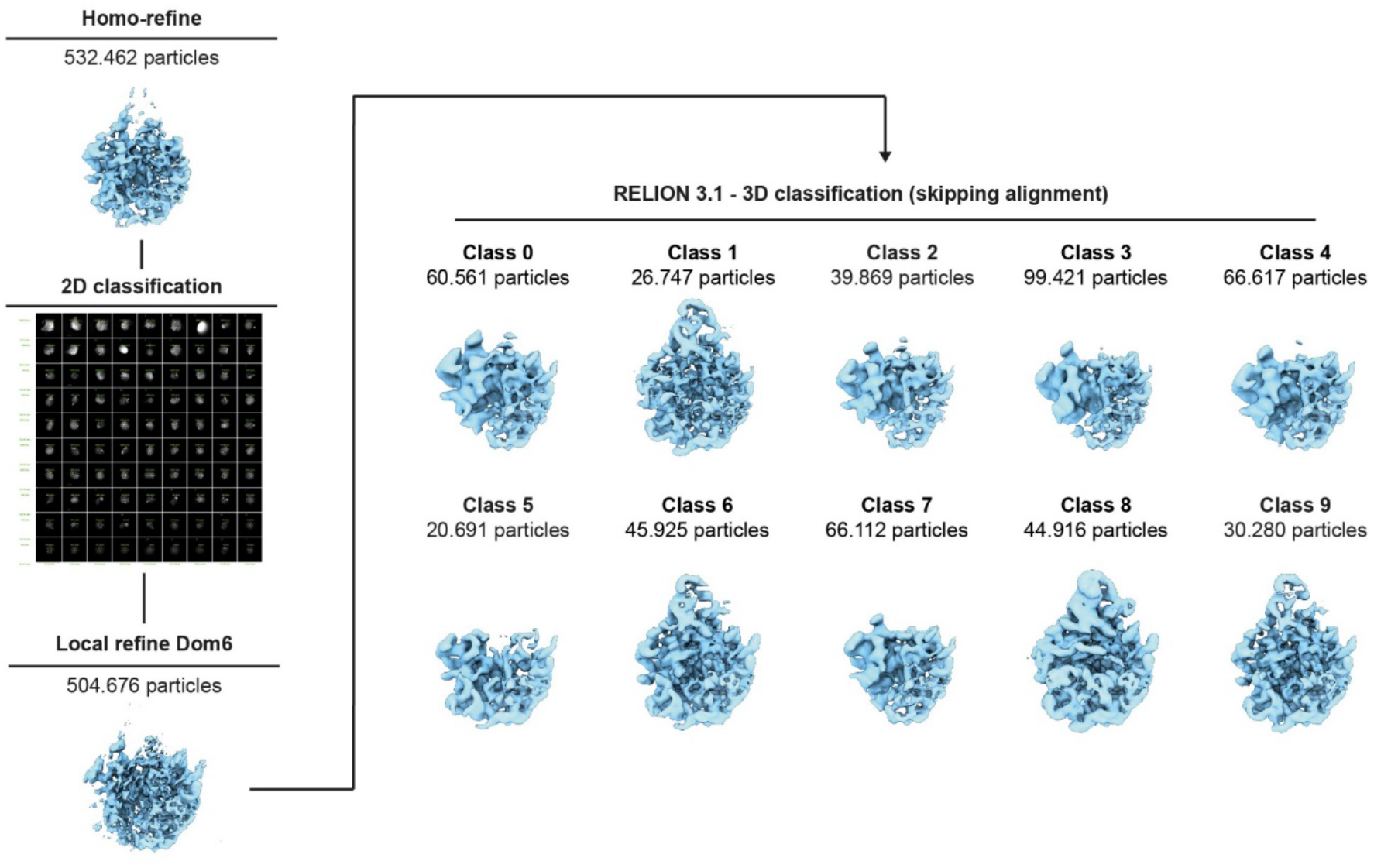
Control sorting scheme. Ribosomal particles from Extended Data Fig.2 were aligned to a consensus 50S map, cleaned by 2D classification and aligned to domain 6 using local refinement. Subsequently, aligned particles were subjected to 3D classification skipping alignment in Relion 3.1.

**Extended Data Fig. 7:**
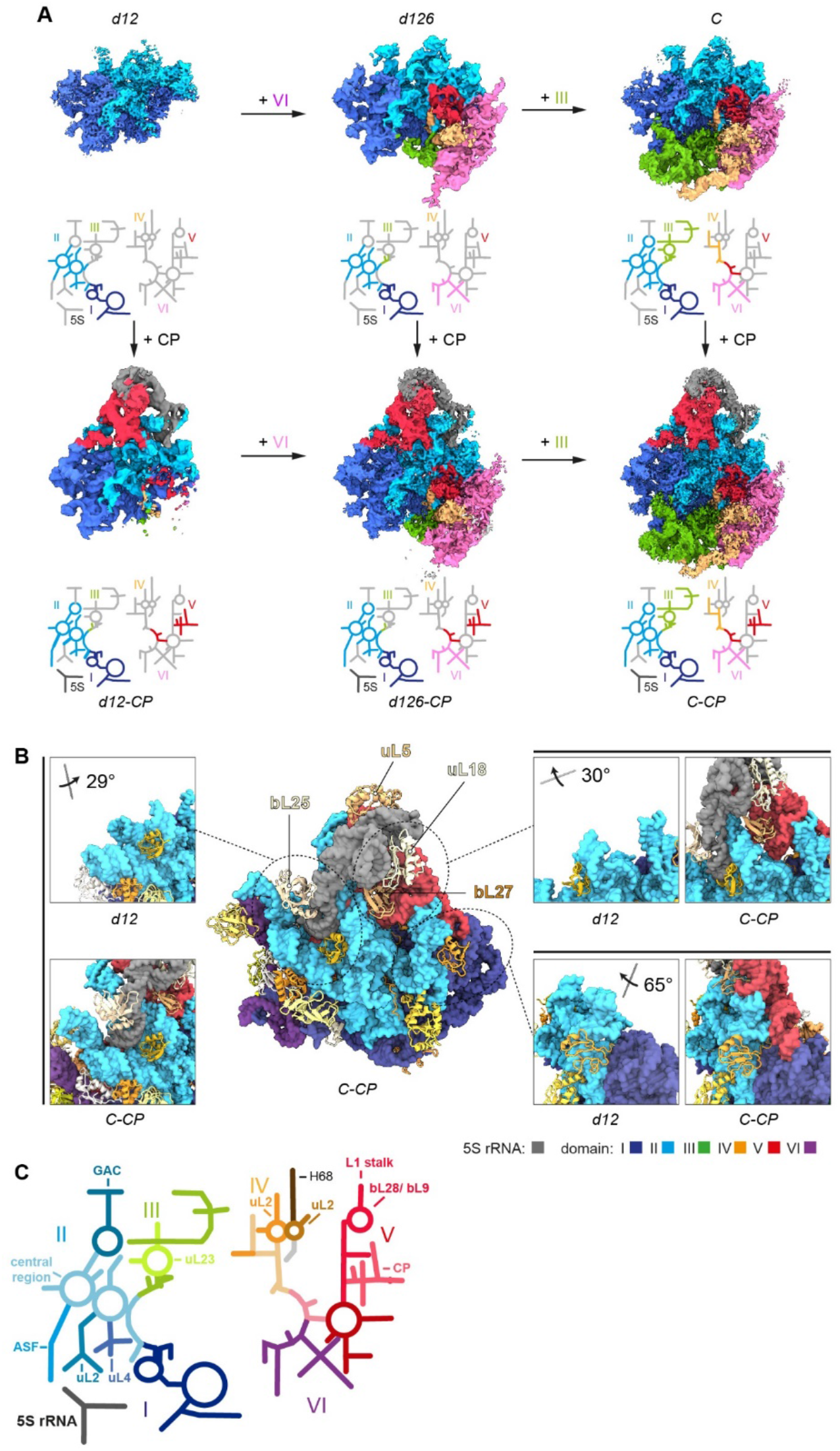
Docking of the 5S RNP. **A)** Cryo-EM maps of states *d12* (*-CP*), *d126* (*-CP*) and *C* (*-CP*), without and with cryo-EM density for the CP, dependent on docking of the 5S RNP, consisting of 5S rRNA, uL5 and uL18. **B)** Models of indicated states with rRNA as surface model (lowpass-filtered to 5 Å resolution), and L-proteins as cartoons. Selected regions are shown in detail, in absence and presence of the CP. Domains of the rRNA color-coded as indicated. Viewing angles shown relative to the full model (*C-CP*). **C)** 23S rRNA 2D map with the individual domains color-coded as indicated, binding sites of L-proteins and features of rRNA regions labeled. GAC, GTPase associated center (H42-44); CP, central protuberance (H80-88) and 5S rRNA; central region of domain II (pale blue); ASF, A-site finger (H38); L1 stalk (H 76-78).

**Extended Data Fig. 8:**
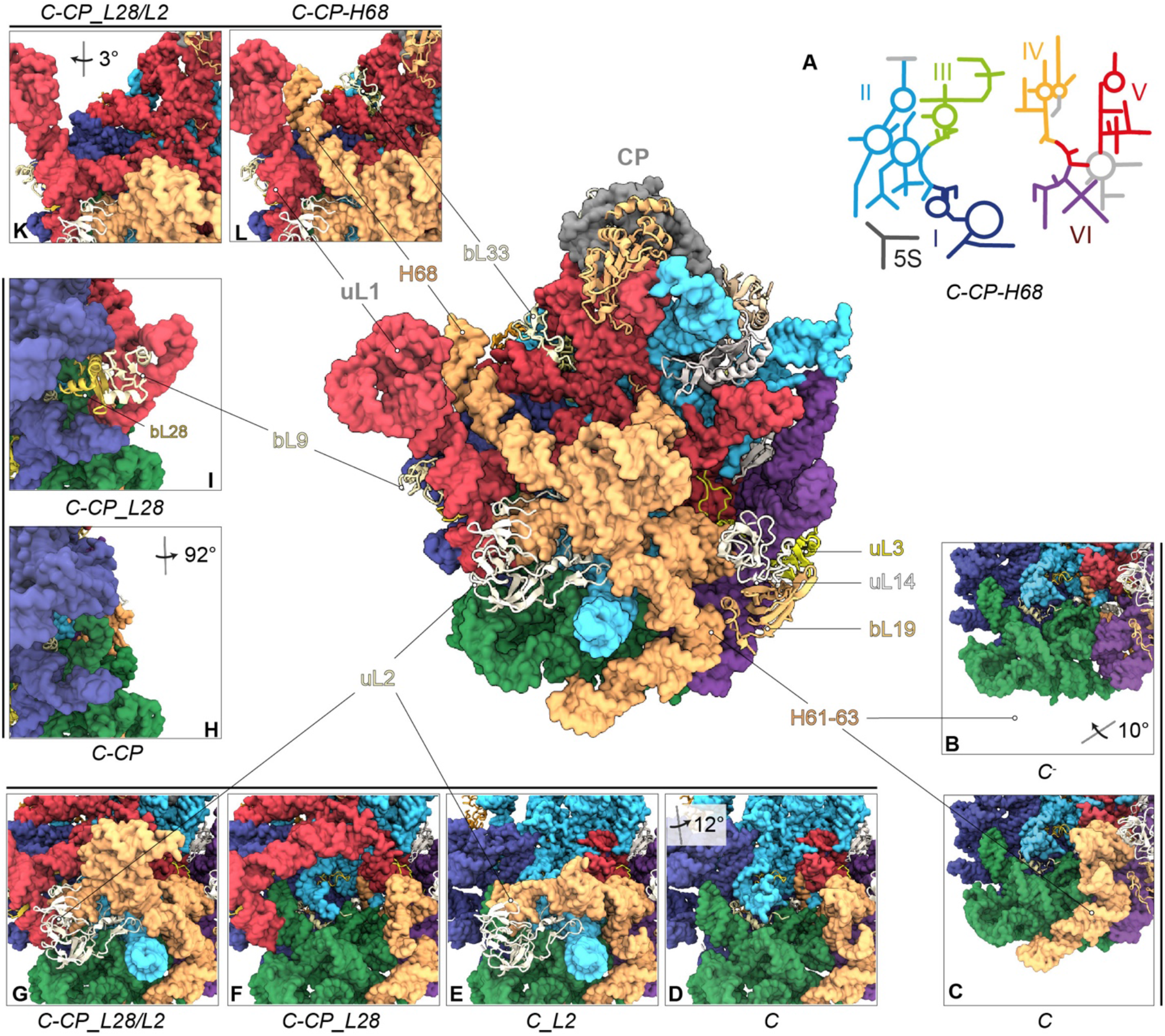
Incremental late 50S assembly. **A)** 2D rRNA map and atomic model of state *C-CP-H68* in crown view. L-proteins appear as cartoons, rRNA as surface model (lowpass-filtered to 5 Å resolution), color-coded as in the 2D-map. Lower sections of states *C*^*-*^ **(B)** and *C* **(C)** lacking or exhibiting helices H61-63, respectively. **D-G)** Lower section of the individual states with absence or presence of uL2. Close-ups of states *C-CP* **(H)** and *C-CP_L28* **(I)**, lacking or exhibiting bL28, bL9 and L1-stalk, respectively. Close-ups of states *C-CP_L28/L2* **(K)** and *C-CP-H68* **(L)** lacking or exhibiting bL33 and H68, respectively. CP, central protuberance. Viewing angles in B), D), H) and K) are shown relative to the full model of *C-CP-H68* (A).

**Movie S1: Relative flexibility of rRNA domains**

For each state, 10 serial Cryo-EM maps along an orthogonal variability trajectory were reconstructed, filtered to 8A and shown as volume series, with the six rRNA domains colored according to the default code. Proteins transparent in gold and silver colors as indicated. Atomic models of proteins shown as ribbons in the same colors. While the domain 1, 3 and 6 are intrinsically rigid, they are flexible relative to each other, which enhances cooperativity. Once formed, the L1 stalk remains flexible, while the CP’s flexibility is terminated by incorporation of bL33, bL35 and uL2.

**Movie S2: Local seeding and cooperativity**

Cryo-EM maps of *d1* states with rRNA domains according to the default color code. Proteins transparent in gold and silver colors as indicated. Atomic models of proteins shown as ribbons in the same colors. Minimal set of L-proteins (uL24, uL22 and uL29) in state *d1* and uL4 and uL23 mediated seeding of adjacent rRNA helices.

**Movie S3: Maturation of the subunit’s core**

Cryo-EM maps of *d1* states with rRNA domains according to the default color code. Proteins transparent in gold and silver colors as indicated. Atomic models of proteins shown as ribbons in the same colors. Minimal set of L-proteins (uL24, uL22 and uL29) in state *d1* and uL4 and uL23 mediated seeding of adjacent rRNA helices.

